# Oxford Nanopore’s 2024 sequencing technology for *Listeria monocytogenes* outbreak detection and source attribution: progress and clone-specific challenges

**DOI:** 10.1101/2024.07.12.603236

**Authors:** Michael Biggel, Nicole Cernela, Jule Anna Horlbog, Roger Stephan

## Abstract

Whole genome sequencing is an essential cornerstone of pathogen surveillance and outbreak detection. Established sequencing technologies are currently challenged by Oxford Nanopore Technologies (ONT), which offers an accessible and cost-effective alternative enabling gap-free assemblies of chromosomes and plasmids. Limited accuracy has hindered its use for investigating pathogen transmission, but recent technology updates have brought significant improvements. To evaluate its readiness for outbreak detection, we selected 78 *Listeria monocytogenes* isolates from diverse lineages or known epidemiological clusters for sequencing with ONT’s V14 Rapid Barcoding Kit and R10.4.1 flow cells. The most accurate of several tested workflows generated assemblies with a median of one error (SNP or indel) per assembly. For 66 isolates, cgMLST profiles from ONT-only assemblies were identical to those generated from Illumina data. Eight assemblies were of lower quality with more than 20 erroneous sites each, primarily caused by methylations at the GAAGAC motif (5′-GAA<u>G6mA</u>C-3 / 3′-GT<u>4mC</u>TTC-5′). This led to inaccurate clustering, failing to group isolates from a persistence-associated clone that carried the responsible restriction-modification system. Out of 50 methylation motifs detected among the 78 isolates, only the GAAGAC motif was linked to substantially increased error rates. Our study shows that most *L. monocytogenes* genomes assembled from ONT-only data are suitable for high-resolution genotyping, but further improvements of chemistries or basecallers are required for reliable routine use in outbreak and food safety investigations.

## Introduction

Whole genome sequencing has become an indispensable tool in genomic epidemiology (Baker et al., 2023; Sintchenko and Holmes, 2015). Currently, most public health and food safety laboratories rely on Illumina short-read sequencing. Oxford Nanopore Technology (ONT) sequencing has emerged as an attractive alternative by offering a cost-effective, accessible, and real-time sequencing approach with long-read data output. Increased error rates, particularly at homopolymer and methylated sites (Delahaye and Nicolas, 2021; Sereika et al., 2022), have so far hindered its use for pathogen tracking, where high accuracies are essential for determining the isolate’s relatedness. However, recent advancements in ONT’s chemistry, flow cells, and basecalling algorithms have significantly improved its accuracy.

*Listeria monocytogenes* is a foodborne bacterial pathogen responsible for high human fatality rates and substantial financial losses in the food industry. Rapid and reliable genotyping is crucial for preventing and controlling outbreaks. *L. monocytogenes* is divided into the four major lineages I, II, III, and IV. Clones relevant to the clinical and food safety context almost exclusively belong to lineages I and II (Moura et al., 2016). Like most bacterial species, *L. monocytogenes* strains commonly carry methyltransferases, often alongside restriction enzymes as part of restriction-modification (RM) systems (Beaulaurier et al., 2019). By cleaving unmethylated DNA, RM systems defend their host against foreign DNA such as phages or plasmids. Because PCR-free ONT strategies sequence native DNA, base modifications such as methylations can be identified but occasionally lead to inaccurate basecalls.

The objective of our study was to determine if ONT sequencing could provide high-quality *L. monocytogenes* assemblies suitable for tracking transmissions. To this end, we ONT-sequenced 78 *L. monocytogenes* with diverse methyltransferases and methylation patterns.

## Methods

### Selection of bacterial isolates

The Swiss National Reference Centre for Enteropathogenic Bacteria and Listeria (NENT) receives and routinely Illumina-sequences clinical *Listeria* isolates from diagnostic laboratories or medical centres in Switzerland. For food safety investigations, isolates from foods and the environment of food-processing facilities are also collected and analyzed. In this study, we selected 78 isolates with available Illumina data from the NENT collection. These included 49 isolates representing the species diversity belonging to the major *L. monocytogenes* lineages I (n = 24), II (n = 53), and III (n = 1). An additional 29 isolates belonged to six cgMLST clusters comprising genetically related (cgMLST distance ≤ 10 alleles) and epidemiologically linked isolates within ST7, ST9, ST121, and ST388. Specifically, the ST7 clone (here represented by 8 facility environment isolates and a clinical blood isolate) persisted in the environment of a poultry meat cutting facility for more than three years and was linked to one bacteremia case. Three related ST9 clones (represented by 3, 3, and 4 facility environment isolates in each cgMLST cluster) persisted in the same meat processing facility for at least one year each. The ST121 clone (here represented by 3 salmon product isolates and two facility environment isolates) was repeatedly recovered from a salmon processing facility over more than two years. The ST388 clone (here represented by 4 clinical blood isolates, a trout product isolate, and a facility environment isolate) was responsible for an outbreak in Switzerland in 2022 linked to smoked trout causing at least 16 bacteremia cases in humans.

### Whole genome sequencing

For Illumina sequencing, DNA was extracted using the DNAeasy Blood and Tissue Kit (Qiagen) or the MagPurix Bacterial DNA Extraction Kit (Zinexts). Libraries were prepared with the Nextera DNA Flex Library Preparation Kit (Illumina) and sequenced using the Illumina MiniSeq platform with 2 × 150-bp paired-end chemistry to a minimum coverage of 50x.

For ONT sequencing, DNA was extracted using the MagPurix Bacterial DNA Extraction Kit (Zinexts). Libraries were prepared using the SQK-RBK114.24 Rapid Barcoding Kit (Oxford Nanopore) according to the “Nanopore-only Microbial Isolate Sequencing Solution” protocol. Libraries were sequenced on MinION devices using R10.4.1 flow cells.

### Bioinformatic analysis

Illumina reads were trimmed and quality filtered using fastp 0.22.0 (Chen et al., 2018). Illumina-only assemblies were generated using SPAdes 3.15.5 (Bankevich et al., 2012) implemented in shovill 1.0.9 (github.com/tseemann/shovill) with a minimum contig length of 200 bp.

For nanopore data, simplex basecalling, demultiplexing, and adapter and barcode trimming were performed with dorado 0.7.0 (github.com/nanoporetech/dorado) in conjunction with the sup@v5.0 (dna_r10.4.1_e8.2_400bps_sup@v5.0.0) model. The models sup@v4.3 and hac@v5.0 were also tested, as indicated. Reads with mean Q-scores of <10 were discarded during basecalling. Nanopore reads were filtered to a minimum read length of 1000 bp and quality controlled using nanoq 0.10.0 (Steinig and Coin, 2022) and subsampled with rasusa 2.0.0 (option --genome-size 3m) (Hall, 2022). ONT-only assemblies were generated from the subsampled reads with flye 2.9.2 (Kolmogorov et al., 2019) and circular contigs rotated using Dnaapler 0.7.0 (Bouras et al., 2024a). Assemblies were then polished with the subsampled reads using flye polisher 2.9.2 (nanohq option, one iteration). Additional polishers that were tested on all assemblies included racon 1.5.0 (Vaser et al., 2017) and medaka 1.12.0 (github.com/nanoporetech/medaka), as specified. Medaka was used in conjunction with the r1041_e82_400bps_sup_v5.0 (referred to as v5.0) or r103_min_high_g360 (referred to as g360) models. Additional correction of selected flye-polished assemblies was attempted by calling variants with Clair3 v1.0.8 (Zheng et al., 2022) (options --include_all_ctgs -- no_phasing_for_fa --haploid_precise, model r1041_e82_400bps_sup_v500) and generating consensus assemblies using bcftools 1.15.1 (Li, 2011). Minimap 2.28-r1209 (Li, 2018) (option map-ont) was used to generate input alignments for racon and Clair3.

Errors in ONT-only assemblies were determined using the compare_assemblies.py script available at github.com/rrwick/Perfect-bacterial-genome-tutorial, utilizing the edlib (Šošić and Šikić, 2017) aligner. Each ONT-only assembly was compared to its corresponding assembly that was polished with Illumina data using Polypolish 0.6.0 (Bouras et al., 2024b). As misassembled plasmids (such as multiple copies in a single contig) compromise downstream analyses, only chromosomal sequences were considered in comparisons. A divergent site was defined as any number of changes (indels or SNPs) within a 30 bp region. Assembly completeness and potential contamination were verified with checkm 1.2.2 (Parks et al., 2015). Sequence types were identified with mlst 2.22.0 (github.com/tseemann/mlst). Core genome (cgMLST) distances were determined using pyMLST 2.1.6 (options -c 0.9 -i 0.9) (Biguenet et al., 2023) in combination with the *L. monocytogenes* cgMLST scheme v2.1 (1701 loci) (Ruppitsch et al., 2015) available at cgmlst.org. Profiles from Illumina-only assemblies were used as references for determining pairwise cgMLST distances. Methyltransferases were identified using abricate (github.com/tseemann/abricate) (minimum coverage / identity 50% / 50%) in combination with the Rebase database (Roberts et al., 2023). To detect methylated bases, ONT raw data was basecalled using dorado 0.7.0 in conjunction with sup@v5.0.0 and the models sup@v5.0.0_4mC_5mC@v1 and sup@v5.0.0_6mA@v1. Methylated motifs were identified with modkit 0.3.1 (github.com/nanoporetech/modkit) (find-motifs option).

## Results

### Highly accurate assemblies obtained after sup-v5.0 basecalling and medaka(g360) polishing

For this study, 78 *L. monocytogenes* isolates with available Illumina data were re-sequenced using the ONT V14 Rapid Barcoding Kit to a minimum depth of 100x. The isolates were selected from diverse lineages, covering 37 distinct sequence types and harbouring various methyltransferases (Supplementary Table 1). A subset of 29 isolates belonged to known cgMLST clusters within ST7, ST9, ST121, or ST388.

We compared different basecalling models, assembly polishers, and sequencing depths to determine their effect on assembly accuracy. Basecalling with the recently released sup@v5.0 model resulted in significantly higher read qualities (median Q-score of >Q10 reads: 23.2) compared to the sup@v4.3 (median 20.7) or hac@v5.0 (median 17.0) models (Figure 1). This improvement in read quality was reflected in the assembly accuracy: the 78 assemblies generated with flye + flye polisher from sup@v5.0 data at a coverage of 100x differed at 0 to 52 chromosomal sites (median: 2.5 sites) from the corresponding assemblies corrected with Illumina data. In assemblies from sup@v4.3 and hac@v5.0 data, the median error rates were more than 2fold and 12fold higher, respectively (Figure 1).

**Figure 1.**
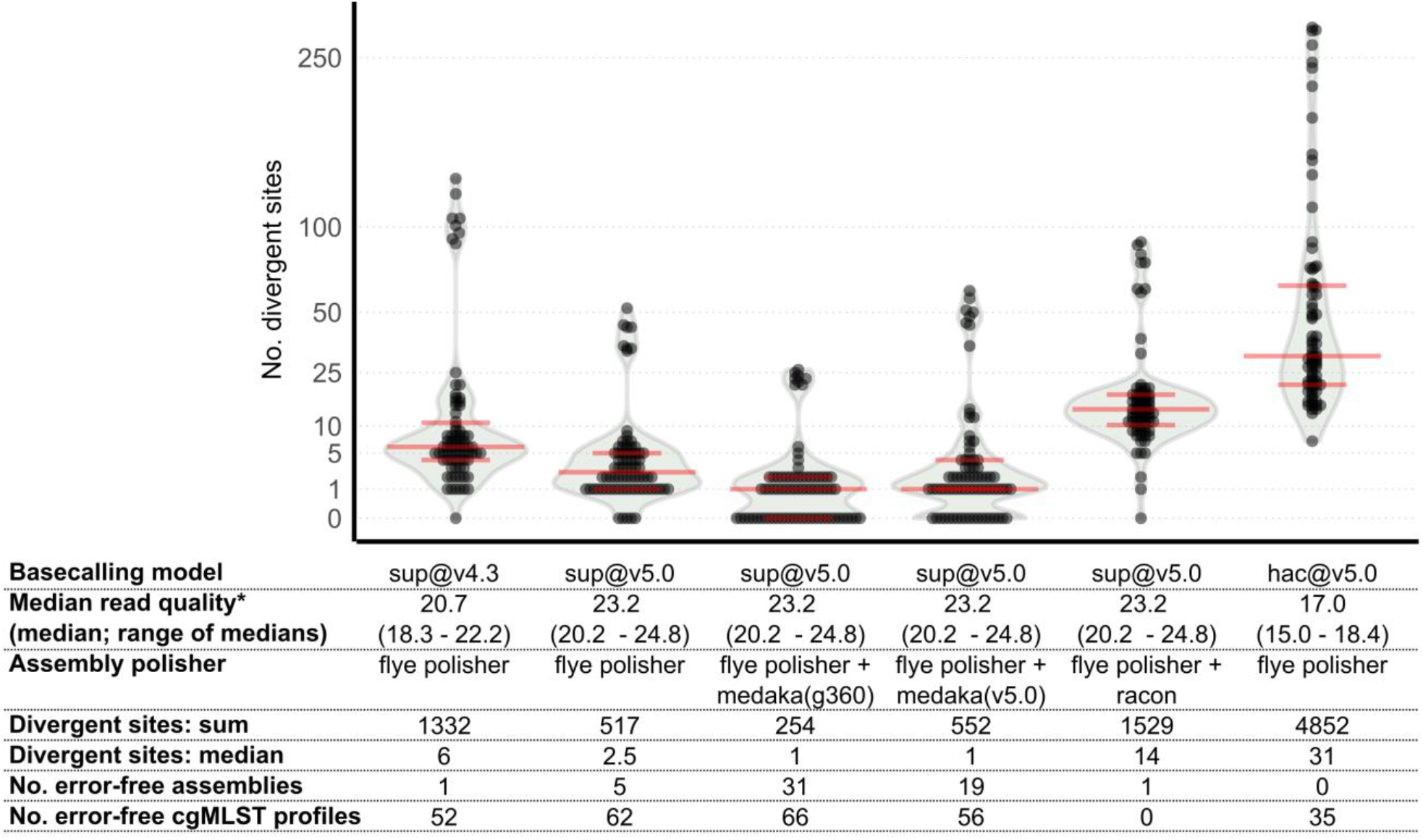
Number of errors in 78 *L. monocytogenes* assemblies generated from ONT-only data at 100x coverage using different basecalling models and polishing approaches. Illumina-polished (hybrid) assemblies were used as references. Only chromosomal contigs were considered in the analysis. The models r103_min_high_g360 (g360) or r1041_e82_400bps_sup_v5.0 (v5.0) were used for medaka polishing as indicated. *Not counting reads with median Q-scores of <10 (discarded during basecalling)

The accuracy of assemblies from sup@v5.0 data could be further improved by additional polishing with medaka in conjunction with the r103_min_high_g360 (g360) model. This approach eliminated about half of the remaining errors (Figure 1). The resulting assemblies had between 0 and 26 erroneous sites (median of 1), and 31 of the 78 assemblies were error-free. By contrast, alternative polishing approaches utilizing medaka with the r1041_e82_400bps_sup_v5.0 (v5.0) model or racon often introduced errors into assemblies that were previously polished with flye (Figure 1).

Assemblies generated from sup@v5.0 data subsampled to lower depths were less accurate. Although at a coverage of 50x, the median error rate remained low (1 error), the eight assemblies with the lowest accuracy had ∼50% more errors compared to the corresponding assemblies generated at 100x coverage (Figure 2).

**Figure 2.**
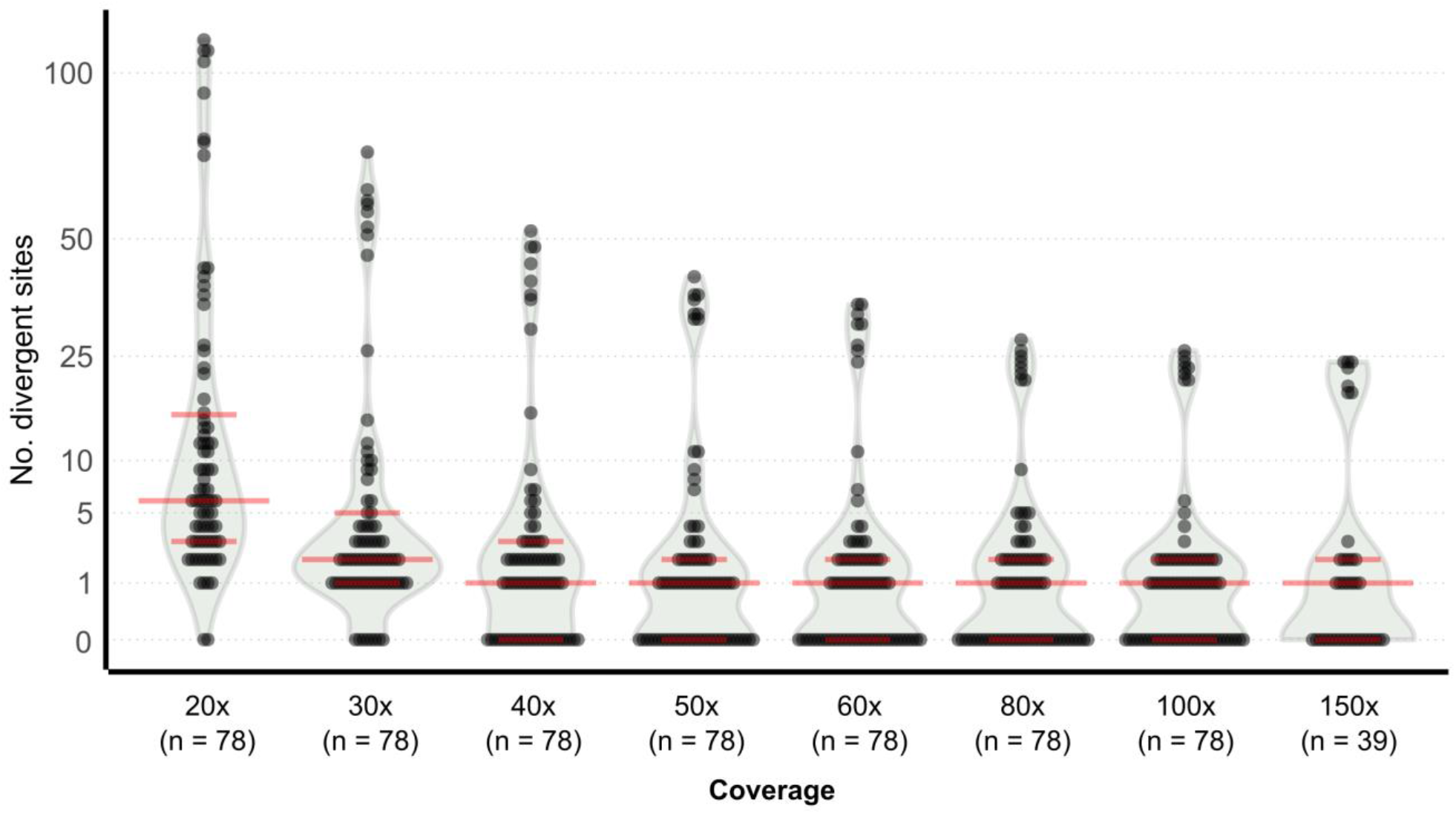
Number of errors in 78 *L. monocytogenes* assemblies generated from ONT-only data subsampled to various sequencing depths. Data were basecalled using the sup@v5.0 model and assemblies were generated with flye followed by polishing with flye polisher and medaka (r103_min_high_g360 model). Illumina-corrected assemblies were used as references. Only chromosomal contigs were considered in the analysis.

### Lower-quality assemblies are associated with GAAGAC-targeting methyltransferases

Of the 78 assemblies generated with the optimized workflow (100x coverage, sup@v5.0 + flye + flye polisher + medaka [g360]), 70 were highly accurate and differed at ≤6 sites from the corresponding Illumina-polished assembly. The remaining eight assemblies had 21 to 26 erroneous sites (Figure 1). In these lower-quality assemblies, most errors (18 to 23 per assembly) occurred at GAAGAC motifs, besides a few errors at GCWGC motifs and homopolymer sites. Low-quality bases at GAAGAC and GCWGC motifs in raw reads were associated with methylations: all eight isolates (belonging to ST3, ST5, and ST121) contained a restriction-modification (RM) system (REBASE: Lmo40025ORF10640P) with two methyltransferases known to target GAAGAC. This system was absent from all higher-quality assemblies. Modified basecalling and analyses with Modkit identified an N^6^-methyladenine (6mA) at this motif (5′-GAAG6mAC-3′) and an N^4^-methylcytosine (4mC) at its opposite strand (3′-GT4mCTTC-5′). Notably, of around 1000 GAAGAC / GTCTTC sites predicted to be methylated in each of these isolates, only ∼2% were incorrectly called. Six of the eight lower-quality assemblies (all from ST121) additionally carried an RM system (REBASE: Lmo2HF33ORFAP) targeting GCWGC (G5mCWGC), which was associated with 2 to 4 additional errors per assembly. Alternative workflows employing either higher sequencing depths (150x), basecalling with the sup@2023-09-22_bacterial-methylation (rerio) model, stricter quality filtering (discarding <Q15 reads), or additional correction of flye-polished assemblies by applying variants called with Clair3 v1.0.8 (Zheng et al., 2022) (r1041_e82_400bps_sup_v500 model) did not improve these lower quality assemblies.

By querying the REBASE database, putative methyltransferases linked to RM systems were detected in overall 75 of the 78 assemblies, with up to five methyltransferases per assembly (Supplementary Table 1). In total, 70 unique methyltransferases were identified, including one (REBASE: M.Lmo0168ORF1995P) that was part of the *mzaABCDE* phage defence system. Modified basecalling predicted highly methylated motifs (at >300 genomic sites) in 64 assemblies (Supplementary Table 1). Among the 50 identified distinct methylation motifs, only methylations at GAAGAC and its complementary strand were linked to substantially increased error rates.

### Effect of assembly errors on cgMLST allele calling and clustering

In pathogen surveillance, potential transmission and persistence clusters are typically investigated at the core genome MLST level. Here, after applying the flye + flye polish + medaka(g360) workflow (100x coverage), 66 out of the 78 ONT-only assemblies produced identical cgMLST profiles as the corresponding Illumina-only assemblies. Four additional assemblies differed by only one or two alleles. The remaining eight assemblies – all lower-quality assemblies from ST3 (n = 1), ST5 (n =1), and ST121 (n = 6) with the GAAGAC-targeting RM system – had 4 to 10 erroneous alleles, which may result in false-negative results in outbreak investigations.

Indeed, five of the ST121 isolates – obtained from salmon products and the environment of a salmon-processing facility – belonged to a cgMLST cluster with allelic distances of 1 to 7 pairwise alleles according to Illumina-only assemblies (Figure 3). In ONT-only assemblies, the corresponding assemblies differed by 5 to 15 alleles. When strictly applying the common threshold of ≤10 allele differences, two of these isolates would have been mistakenly classified as outside the cluster using ONT-only data.

**Figure 3:**
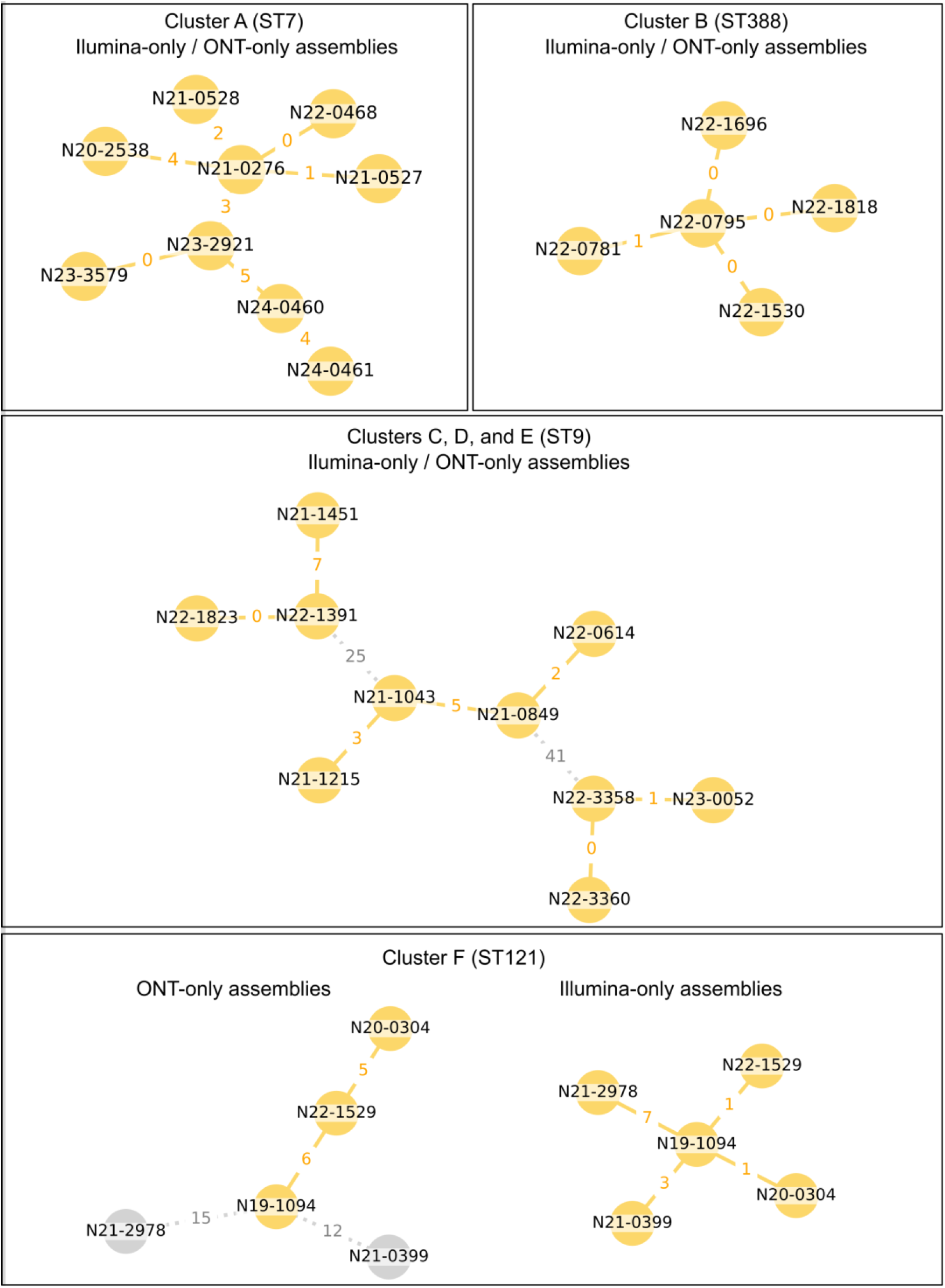
Comparisons of minimum spanning trees based on cgMLST profiles of assemblies generated from Illumina-only versus ONT-only sequencing data. The six *L. monocytogenes* clones within ST7, ST9 (n = 3), ST121, and ST388 are linked to outbreaks or persisted in food processing facilities. Numbers between nodes indicate the allelic distance. Isolate nodes and edges assigned to cgMLST clusters (distance ≤10 alleles) are coloured yellow. For assemblies from isolates within the ST7, ST9, and ST388 clusters, Illumina-only and ONT-only workflows produced identical cgMLST profiles. ONT-only assemblies from the ST121 cluster contained multiple mismatching alleles, resulting in false-negative clustering.

Five additional cgMLST clusters belonging to ST7, ST9 (3 clusters), and ST388 and linked to outbreaks or persistence in food processing facilities (see methods) were investigated. For all 24 isolates from these clusters, cgMLST profiles from ONT-only assemblies were identical to those from the corresponding Illumina-only assemblies, hence resulting in identical pairwise distances between isolates and accurate clustering (Figure 3).

## Discussion

Genomic epidemiology investigations require high sequencing accuracies approaching 100% at the consensus level to effectively discriminate between closely related strains. Our analysis of 78 *L. monocytogenes* isolates showed that – after employing sup@v5.0 basecalling and assembly polishing with flye and a specific medaka model – most ONT-only assemblies were highly accurate and suitable for high-resolution genotyping. However, in clones within *L. monocytogenes* ST3, ST5, and ST121 that carried a specific RM system, methylated GAAGAC / GTCTTC sites were often miscalled and produced unreliable genotyping results at both wgSNP and cgMLST levels. When applied to a real-world case of an ST121 clone persisting in a food-processing environment and carrying this system, the isolates failed to cluster at the widely used threshold of ≤10 differing alleles, unlike corresponding Illumina-only assemblies. Notably, ST3, ST5, and ST121 – each encompassing subclones carrying this RM system – are among the dominant *L. monocytogenes* lineages recovered from food and clinical sources (Coipan et al., 2023; Maury et al., 2016). Overall, the 78 assemblies contained diverse methyltransferases and 50 frequently methylated motifs. Among those, GAAGAC / GTCTTC was the only motif with distinct methylation types (6mA and 4mC) on opposite strands (i.e. 5′-GAAG6mAC-3′ / 3′-GT4mCTTC-5), possibly presenting a challenge for assembly and polishing algorithms.

Several previous studies evaluated the performance of ONT for genotyping pathogens. For instance, Hong et al. (Hong et al., 2024) reported fewer than 10 mismatching cgMLST alleles compared to Illumina-only assemblies for 12 *L. monocytogenes* and 23 *Salmonella enterica* isolates (using the RBK114.24 kit, R10.4.1 flow cells, and dorado 0.5.0 with the sup@4.3 model). In a study on 356 multi-drug resistant isolates, Landmann et al. (Landman et al., 2024) described highly congruent wgMLST profiles (0 to 9 alleles difference) for ONT-only and Illumina-only assemblies from *K. pneumoniae, E. coli, E. cloacae* complex, *C. freundii, A. baumannii* complex and MRSA, but noted lower accuracies for *P. aeruginosa* (using RBK114.24, R10.4.1 flow cells, dorado v0.3.2 with sup@2023-09-22_bacterial-methylation). Lohde et al. (Lohde et al., 2023) reported discrepancies in cgMLST distances that prevented accurate clustering for multiple *K. pneumoniae* isolates due to suspected methylations (using V12 / V14 NBD kits, R10.4 / R10.4.1 flow cells, and guppy v6.4.6 / dorado v0.3.0 with various models).

Methylation-associated errors observed in our study may be prevented or reduced by employing alternative library preparation kits or bioinformatic workflows, though these approaches have their limitations. Whole genome amplification-based approaches replace methylated bases during amplification but are more costly due to lower sequencing data yields and often result in fragmented assemblies due to shorter read lengths (Chiou et al., 2023; Lohde et al., 2023). ONT duplex sequencing improves read and assembly quality (Hall et al., 2024) but for high duplex depth currently requires ONT’s ligation kits which are less suitable for high-throughput bacterial sequencing. The software modpolish uses a reference-based strategy to correct assembly errors but relies on closely related and high-quality reference genomes, thereby introducing bias (Chiou et al., 2023). Finally, the masking of ambiguous bases was shown to effectively reduce errors but compromise downstream analyses such as allele calling for cgMLST (Lohde et al., 2023).

In conclusion, this study shows that ONT sequencing facilitates highly accurate assemblies but is not yet consistently reliable for investigating bacterial outbreaks and contamination sources. We expect that upcoming releases of improved basecallers will resolve the remaining issues and enable accurate genotyping of all clones.

